# Bemovi, software for extracting Behaviour and Morphology from Videos

**DOI:** 10.1101/011072

**Authors:** Frank Pennekamp, Nicolas Schtickzelle, Owen L. Petchey

## Abstract

1. Microbes are critical components of ecosystems and vital to the services they provide. The essential role of microbes is due to high levels of functional diversity, which are, however, not always mirrored in morphological differentiation hampering their taxonomic identification. In addition, the small size of microbes hinders the measurement of morphological and behavioural traits at the individual level, as well as interactions between individuals.
2. Recent advances in microbial community genetics and genomics, flow cytometry and digital image analysis are promising approaches, however they miss out on a very important aspect of populations and communities: the behaviour of individuals. Video analysis complements these methods by providing in addition to abundance and trait measurements, detailed behavioural information, capturing dynamic processes such as movement, and hence has the potential to describe the interactions between individuals.
3. We introduce bemovi, a package using R - the statistical computing environment - and the free image analysis software ImageJ. Bemovi is an automated digital video processing and analysis work flow to extract abundance and morphological and movement data for numerous individuals on a video, hence characterizing a population or community by multiple traits. Through a set of functions, bemovi identifies individuals present in a video and reconstruct their movement trajectories through space and time, merges measurements from all treated videos into a single database to which information on experimental conditions is added, readily available for further analysis in R.
4. We illustrate the validity, precision and accuracy of the method for experimental multi-species communities of protists in aquatic microcosms. We show the high correspondence between manual and automatic counts of individuals and illustrate how simultaneous time series of abundance, morphology and behaviour are constructed. We demonstrate how the data from videos can be used in combination with supervised machine learning algorithms to automatically classify individuals according to the species they belong to, and that information on movement behaviour can substantially improve the predictive ability and helps to distinguish morphologically similar species. In principle, bemovi should be able to extract from videos information about other types of organism, including “microbes”, so long as the individuals move relatively fast compared to their background.

## 1 Introduction

Microbes are crucial components of all ecosystems providing important services such as organic matter decomposition, production of biomass and oxygen, or carbon storage (Kirchman 2012). Nevertheless, they are still mostly treated as a ‘blackbox’ in terms of their phenotypes, ecology and behaviour (Finlay 2004). This is mainly due to methodological limitations which can describe microbes at best at the population or community level. While these approaches provided important insights into the functional diversity of microbes, limitations in describing microbes at the individual level are in contrast with the insight that most ecological and evolutionary processes are driven by individuals (Bolnick *et al.* 2003). The same limitations apply to the use of microbial model systems such as protists, which have a long and successful tradition in testing ecological and evolutionary theory (Gause 1934, Holyoak & Lawler 2005, Elena & Lenski 2003, Kawecki *et al.* 2012).

Advances in microbiology have been driven by technological developments ever since Antonie van Leeuwenhoek invented the compound microscope (Kreft *et al.* 2013). New technologies like metagenomic studies, flow cytometry, digital image analysis and single-cell microbiology provide insights about the structure and composition of microbial communities, as well as the morphology, function and ecology of microbes at the individual level (Kreft *et al.* 2013, Brehm-Stecher & Johnson 2004). Whereas these new technologies are powerful in showing differences among individuals in physiology or morphology, they miss out on an important component of the individual phenotype: behaviour, which is an important factor shaping interactions among species (McGill & Mittelbach 2006). Digital video analysis can adequately complement these approaches by providing quantitative descriptions of behaviour via automated tracking (Dell *et al.* 2014), which is collected at the individual level in addition to abundance and morphological data. Such methods apply to all types of empirical systems where individual members of the community are characterized by movement, e.g. they should work on samples freshly taken in the natural environment and also for the widely used micro- and mesocosm systems.

There are now different software to perform tracking, but these are focused on extracting movement and behavioural information only, without associated morphology and abundance data. They are also stand-alone, sometimes lack efficient ways of dealing with large numbers of files, and may be difficult to customize and automate. Dell *et al.* (2014) give an extensive overview of these software and their strengths and weaknesses. While previous demonstrations relying on digital image analysis (e.g. Pennekamp & Schtickzelle 2013, Mallard *et al.* 2013) tested and validated work flows aimed at single-species microcosms, extending the capability of such systems to larger communities is required (Gaston & O’Neill 2004). Some success with automatic classification of species was achieved with protists in activated wastewater sludge (Amaral *et al.* 2008) and automated systems to analyse abundance and quantify trait distributions during large-scale marine monitoring schemes (e.g. Zoo/PhytoImage (http://www.sciviews.org/zooimage/) (Bell & Hopcroft 2008)). These show that such efforts are worthwhile even with challenging field-collected samples. No studies so far however used automated video analysis, which captures the dynamic movement behaviour of the study organisms and therefore potentially a characteristic signature for species identities.

Automated video analysis usually consists of three main steps: video acquisition, video processing/analysis, and data interpretation (Dell *et al.* 2014). To fulfil the latter two steps, we introduce a new R package, bemovi, and show its validity and scope of application. For guidance on the image acquisition setup, refer to Dell *et al.* (2014) or Pennekamp & Schtickzelle (2013). Bemovi is an automated digital video processing and analysis work flow to extract abundance, morphological and movement data for numerous individuals on a video, hence characterizing a microbial population or community by multiple traits. We illustrate how the individual trait data can be used to predict species identity in a multi-species community, and how the characteristics of the movement improve the predictive ability of the classification model compared to morphological data only. We then derive population abundance by counting the individuals of each species and validate these against manual counts taken simultaneously for both single and multi-species communities.

## 2 Description of the bemovi package and its functions

**Table 1:** Overview of functions provided with the bemovi package. Functions are ordered according to the analysis flow.

| Step | function name | short description |
| --- | --- | --- |
| Setup | check_video_file_names<br>check_threshold_values | checks whether video files are either of *.avi or *.cxd format<br>assists finding manually an appropriate threshold for image segmentation |
| Identify and track particles | locate_and_measure_particles<br>link_particles | segments and thresholds video, then runs particle analysis to locate and measure the morphological properties of each particle on all frames<br>reconstructs the movement trajectory of a given particle by linking its coordinates through time and calculates movement metrics such as turning angles and move lengths |
| Merge data | merge_data | creates a database merging morphology and trajectory information with a description of the experimental design |
| Process data | summarize_trajectories<br>filter_data | summarizes the mean morphology and movement and its variability on the trajectory level<br>filters the data, excluding very short, almost non-moving and low detection trajectories |
| Validation | create_overlays | creates an overlay of the original video and the trajectories identified in the segmentation and tracking steps; two different visualization options are possible; if the species was predicted, trajectories can be colored according to the species they were predicted to belong to |

The bemovi work flow relies on two freely available, open source and cross-platform software widely used in the scientific community: R - the statistical computing environment (R Development Core Team 2012) and ImageJ, a powerful image processing and analysis software (Ferreira & Rasband 2010). ImageJ shows considerably better performance than a native R solution for the video processing steps (Pennekamp & Schtickzelle 2013). We therefore built bemovi as a set of modular R functions (Table 1) calling ImageJ and reading its output, creating a seamless work flow that deals efficiently with large numbers of video files and merge results into databases for easy analysis. Additional helper functions are provided to help in setting up and validating the use of bemovi on the experimenter system. bemovi, is readily available from github (https://github.com/pennekampster/bemovi), a solution allowing for easy package upgrading and diffusion, and has been thoroughly tested on Macintosh OS X, Windows 7 & 8 and Ubuntu Linux.

bemovi is built to process a directory containing a set of videos files shot with identical settings, with three main steps (Table 1): (1) identify individuals present in a video and reconstruct their movement trajectories, (2) merge measurements from all treated videos into a single database to which information on experimental conditions is added, (3) perform basic analyses.

In the first step (i.e. identify and track particles), each single video is split into a stack of images (= frames) ordered in time (Dell *et al.* 2014), and each of these images is treated sequentially to locate individuals (function: `locate_and_measure_particles`) and reconstruct their movement trajectories (function `link_particles`). bemovi first uses a dynamic ‘difference image’ segmentation to discriminate individuals from the background, where an image at a constant time offset gets subtracted from the frame to be analyzed (Pennekamp & Schtickzelle 2013). The resulting difference image, only containing the particles that moved, is then binarized (i.e. converted to black and white) using a user defined threshold. Both the time offset and the threshold must be carefully adjusted and validated by the user to minimise segmentation errors (e.g. nearly immobile individuals are erased in the difference image and considered background if the offset is too short). A helper function assists with finding the appropriate threshold (function: `check_threshold_values`). Binarized images are then analysed by the ParticleAnalyzer function of ImageJ, which extracts for each particle, X- and Y-position and morphology (area, mean, minimum and maximum of the grey value, perimeter, width, length and angle with the dominant-axis of a fitted ellipse, circularity, aspect ratio, roundness and solidity) (Ferreira & Rasband 2010).

Once particles have been identified and localised on each frame, their movement trajectories are reconstructed by linking the position of each particle through the stack of images. We use the MOSAIC ParticleTracker plug-in for ImageJ (Sbalzarini & Koumoutsakos 2005) because it can track simultaneously hundreds of particles in an unrestricted viewing field (e.g. cells swim in and out), even when some of them miss out on certain frames due to detection problems. The algorithm deals with occlusions (i.e. when a particle collides with another particle [individual or debris]) conservatively by interrupting the trajectory and starting a new one. For computational efficiency and to avoid the common problem that large variation in size and shape hampers efficient detection in tracking applications (Dell *et al.* 2014), we pass the X- and Y-positions extracted by the ParticleAnalyzer to the ParticleTracker. Two arguments are required for this step: the maximum displacement of particles between two successive frames and the number of frames over which a particle can be linked if missing on some intervening frame(s). They must also be carefully validated to avoid errors (e.g. creating an erroneous link between different particles if displacement and/or link range are too large, or broken links if they are too small). After trajectories are re-constructed, movement metrics are computed for each: move length, absolute angle, turning angle, net squared displacement and gross displacement for the trajectory in its whole. For a detailed description on the calculation and interpretation of these metrics, refer to textbooks on the quantitative analysis of trajectories such as Turchin (1998).

The second step (i.e. data merging) combines the morphology and movement metrics acquired on each video into a single database and links this information with a video description file containing any relevant information on experimental conditions for each video (e.g. treatment level, video capture settings), using the video file name for merging (function: `merge_data`).

In the third step (i.e. data processing), aggregation of the morphology and movement metrics (mean and STD) is performed on the trajectory level (each trajectory has a unique ID of file name and trajectory number). As a final processing step, data can be filtered based on movement and morphology (function: `filter_data`). For error checking and validation of results, a helper function for visualizing the output of bemovi was developed (function: `create_overlays`). The function overlays the original video data with the extracted trajectories and therefore can help to troubleshoot erroneous and incomplete tracking results.

## 3 An illustration on aquatic microbial communities: species classification, population dynamics, and trait dynamics

Video analysis by bemovi provides quantitative measures of the abundance of individuals as well as detailed information on morphology and movement at the individual level. We here illustrate its application to the study of microbial populations and communities of multiple ciliate species in aquatic microcosms using a similar setup as Petchey (2000).

**Figure 1:**
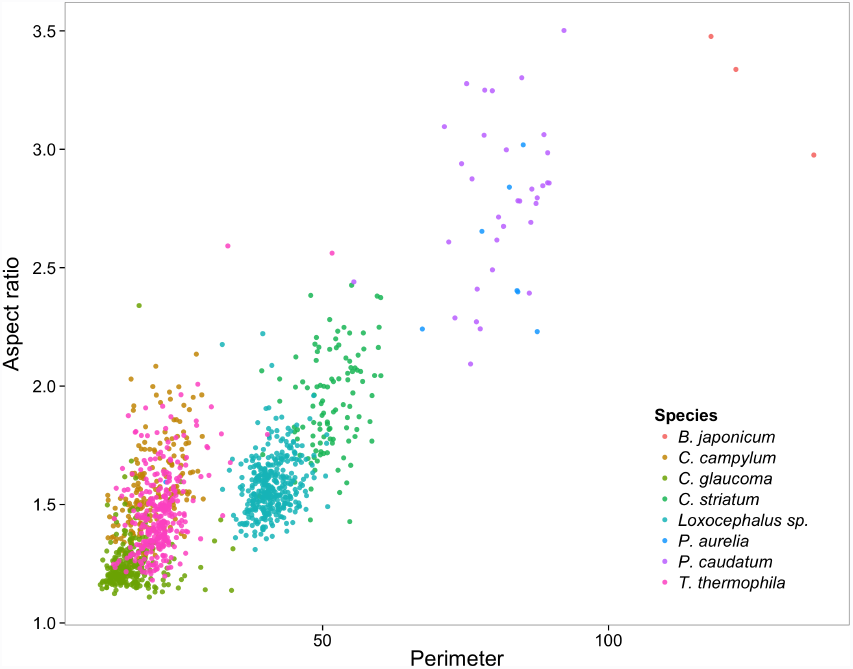
The eight ciliate species’ morphological characteristics, cell perimeter and the aspect ratio (major axis/minor axis of a fitted ellipse). Some species, e.g., *P. caudatum* and *P. aurelia*, show considerable overlap, whereas many species occupy quite distinct areas of the morphological trait space.

**Figure 2:**
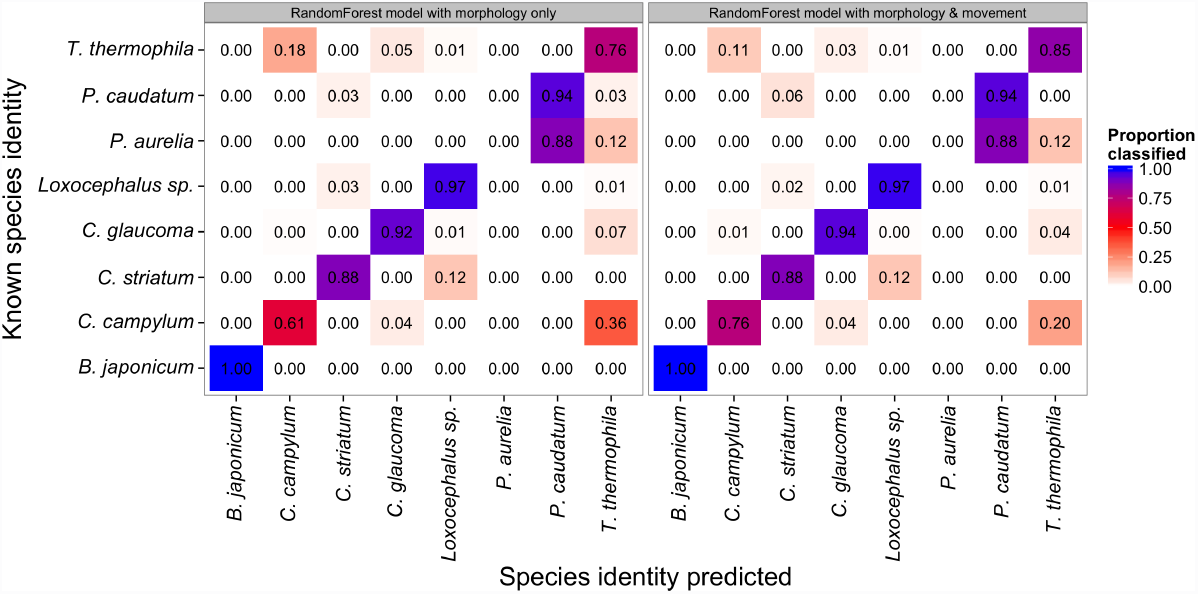
Confusion matrix between eight ciliate species to illustrate the improved classification success (> 89%) when both morphology and movement features are considered, compared to morphology only. Including the movement behaviour decreases the classification error of the randomForest model from 15 to 10% (redish colour off the diagonal). Classification error of the morphologically similar *C. campylum* and *Tetrahymena* (and vice versa) decreases from 36 to 20% and 18 to 11%, respectively. However, highly similar species like *P. caudatum* and *P. aurelia* cannot reliably distinguished as all *P. aurelia* are classified as *P. caudatum.*

Information on traits exhibited by individuals in a mixed community is usually used to predict their species identity as long as sufficient differences in trait space exist (Gorsky *et al.* 2010). Figure 1 shows the differences in two morphological traits (cell size and aspect ratio) among 8 species grown separately in monocultures. Some species like *Paramecium caudatum* and *Paramecium aurelia* show considerable overlap, whereas many species occupy quite distinct areas of the morphological trait space, aiding in their automatic classification. With a supervised machine learning algorithm, the randomForest classification (Breiman 2001, Cutler *et al.* 2007), we were able to reach a high classification success (89%; Figure 2) into the right species classification based on all traits, even if confusion between morphologically similar species was higher (e.g. *Colpidium campylum* from *Tetrahymena thermophila* (and vice versa) 36% and 18%). When the movement characteristics were considered, in addition to morphology, classification error dropped importantly. Highly similar species like *P. caudatum* and *P. aurelia* however, remain indistinguishable.

**Figure 3:**
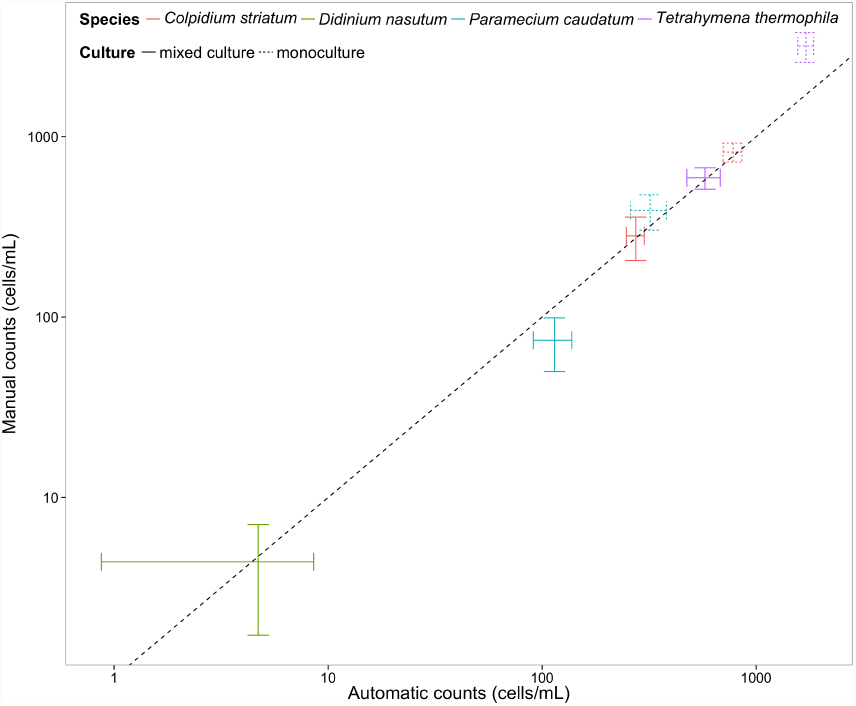
Comparison of manual and automatic estimates of population abundance in single (dashed bars) and mixed species cultures (solid bars). The error bars are +/- 1 STD. The dashed line indicates the 1:1 line.

In another experiment, four ciliate species (*T. thermophila*, *P. caudatum*, *Colpidium striatum* and *Didinium nasutum*) were grown in mono- and mixed cultures, with the aim to compare automatic (using bemovi) and manual counts. We separately took 10 samples from each culture to assess the abundance measurement error, i.e. variability when repeatedly sampling from the same microcosm. Automatic and manual counts showed high correspondence (R^2^: 0.91) (Figure 3), regardless whether counts were performed on mono- or mixed cultures. Measurement error was overall similar between the two methods, though manual counts tend to be more precise at low densities (i.e. *Didinium*), and automatic counts more precise at high densities (i.e. *Tetrahymena*) (Figure 3). These differences are explained by the small and fixed sampling volume (0.0722 mL) used for the automatic counts in this experiment, compared to the variable volume sampled during manual counting (up to 0.5 mL for low density populations, and down to 0.0185 mL for high density cultures. Reducing sampling error in the automated counts could be achieved by increasing the volume videoed. Because the four species used in this experiment are well separated in trait space, very low classification error of about 2% was observed.

**Figure 4:**
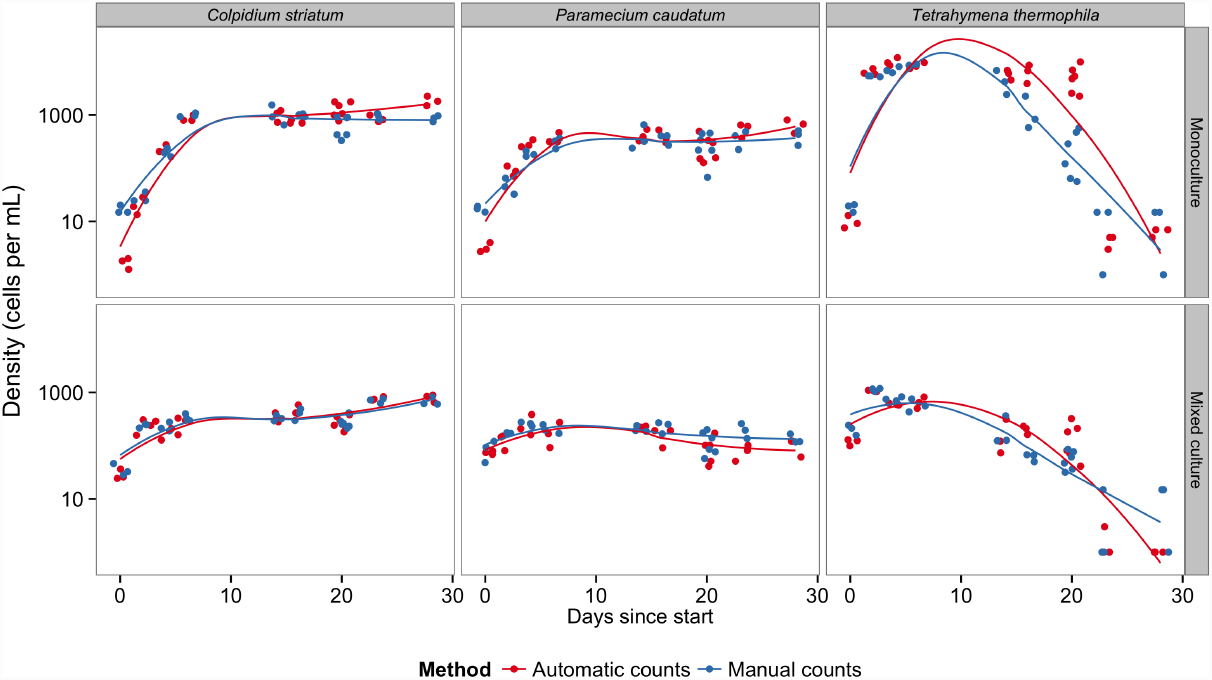
Population dynamics given by manual and automatic counting. Point measures are jittered for easier visualization.

**Figure 5:**
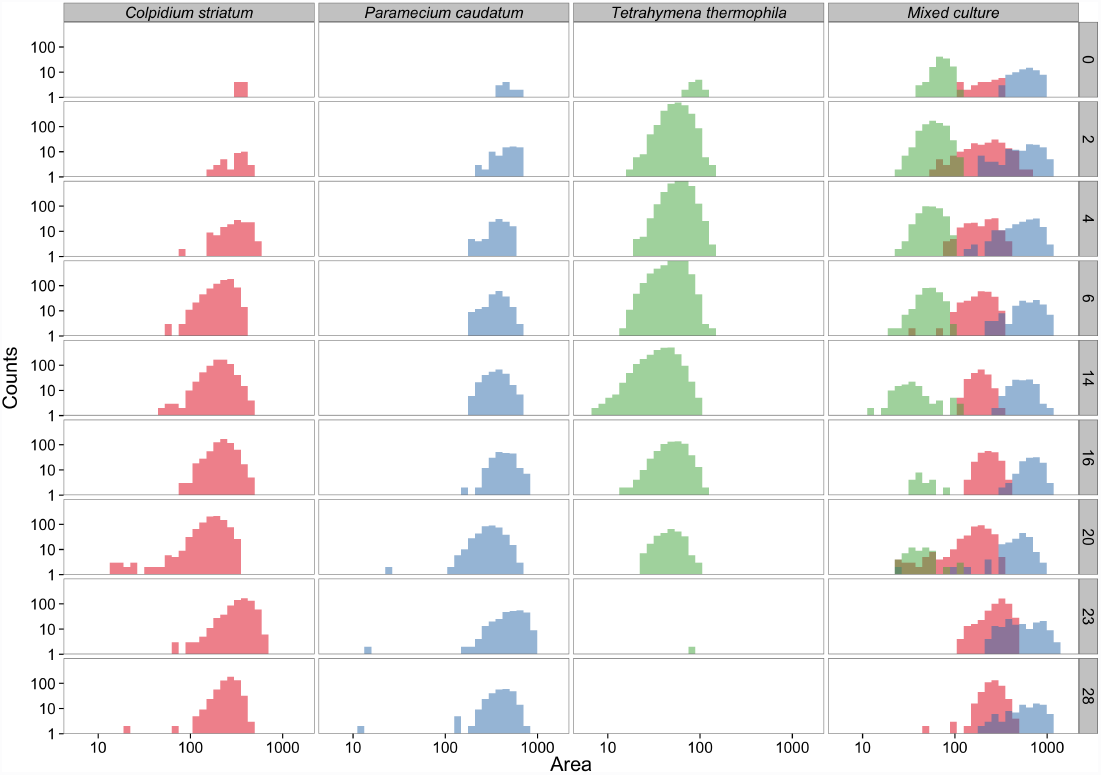
Body size changes of the single species and community during the experiment. At day 20, the medium was partly replaced by fresh medium perturbing the populations and communities. Both axis are on log_10_ scaling.

**Figure 6:**
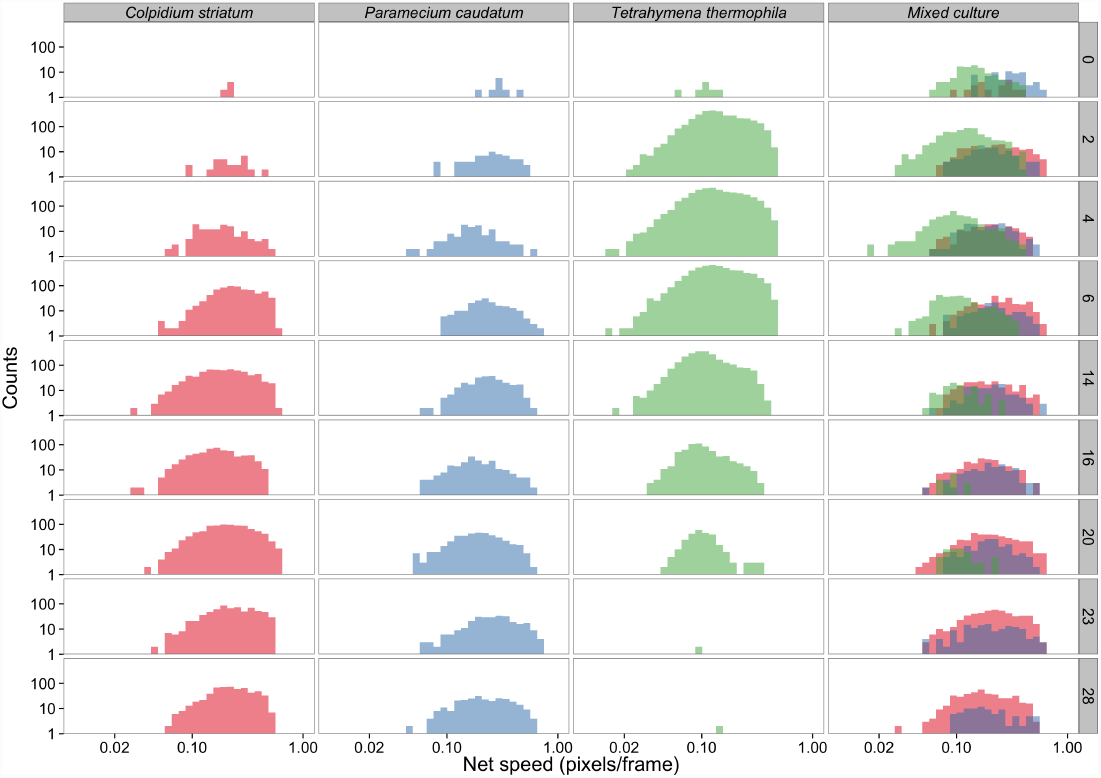
Changes in movement speed in the single species and community during the experiment. At day 20, the medium was partly replaced by fresh medium perturbing the populations and communities. Both axis are on log_10_ scaling.

Finally, we followed mono- and mixed cultures for a period of 28 days and obtained species identification and estimation of abundance both automatically and manually. Both methods captured very well the monoculture growth dynamics of *P. caudatum*, *C. striatum* and *T. thermophila* (Figure 4). Although *T. thermophila* showed some discrepancy between the methods, overall both were closely correlated (Pearson correlation coefficient: 0.93). Furthermore, cell size dynamics (Figure 5) illustrate the ability of bemovi to capture trait dynamics. Over the first twenty days, *C. striatum* and *T. thermophila* decreased in size, whereas *P. caudatum* remained rather stable. After a resource pulse (replacement of 50% of the medium) on day 20, cell sizes increased where present. Figure 6 shows how a dynamic trait such as movement speed changes during the 28 day experiment. The species show stronger overlap in movement speed than in size, but the distribution of speed seems more stable through time and unaffected by the resource pulse on day 20. Such patterns demonstrate the potential of bemovi in trait-based community ecology, though deep analysis of the patterns illustrated here are beyond the scope of this article.

### 3.1 Strengths and limitations

Bemovi shares the advantages of other automated image analysis systems: 1) results (videos) can be stored for later analysis or re-analysis, 2) use of a computer reduces observer bias, 3) video acquisition is usually faster than manual counts and the effort is constant regardless of community complexity (whereas manually counting complex communities can be very time consuming) (Pennekamp & Schtickzelle 2013). For illustration, an experiment including all pairwise combinations of 6 species required 9 person hours to count a total of 108 microcosms. In comparison, video acquisition was achieved in about 2.5 hours. Besides the considerable increase in speed, the work flow presented here extracts far more information than counts because multiple traits are collected simultaneously, at no extra cost. Automatic classification of species also allows to quantify classification uncertainty in multiple species communities in contrast to manual observations. Classifications below a certain threshold (e.g. 90%) could be identified as ‘uncertain’ or measures of classification error integrated in the statistical inference framework.

Some limitations need to considered when using bemovi. First, the tracking of individuals is done in two dimensions, although the environment may be often rather three dimensional. Whereas this may be largely representative for organisms living on a plane (such as ground-dwelling insects for instance), for others such as aquatic organisms this simplification may be problematic for extrapolating the measured movement in the 3D environment. Several studies however show how 2D tracking successfully predicts spread rates even in higher dimensional systems (Giometto *et al.* 2013, Pennekamp 2014). Complex environments with many physical obstacles (e.g. debris particles in our medium) prevent reliable tracking in three dimensions, because individuals may be frequently invisible to one of the three required cameras. Nevertheless, software and hardware rapidly develop ultimately leading to systems performing 3D tracking (Dell *et al.* 2014).

Another limitation, which applies to video tracking in general, is the occurrence of occlusions (i.e. when a particle collides with another particle [individual or debris]) (Dell *et al.* 2014). This can result in errors and their propagation when identifying individuals is required (e.g. in studies of collective movement and social interactions in a swarm of animals). Some recently developed tracking algorithms such as the idTracker (Pérez-Escudero *et al.* 2014) or the Ctrax software (Branson *et al.* 2009) can deal with such situations using powerful ‘fingerprinting’ techniques to keep track of individuals or including probabilistic models, which predict the position after the occlusion, and therefore may be able to maintain the individual identity. Due to the large numbers of individuals tracked simultaneously, the use of computationally intensive ‘fingerprinting’ or manual verification are currently not suited to the goals of bemovi. Given that the work flow is highly modular, another tracking software could easily replace the plug-in used and therefore extend for 3D tracking or having more power in maintaining individual identities.

## 4 Conclusions

Microbes are important for all the ecosystems on the planet, but are still mostly studied at the population or community level. This contrasts with the notion that most ecological processes are driven by individuals (Dell *et al.* 2014). Recent technological advances allow now to study microbes at the individual level and discover whether variation among individuals is as important for microbes as for animals and plants (Kreft *et al.* 2013). The video analysis work flow we presented here complements techniques such as flow cytometry and metagenetic approaches, because it allows to study the behaviour of microbes within populations and communities *in situ*. Characterizing behaviour may thus reveal unrecognised functional diversity, previously masked by low morphological and potentially genetic differentiation.

Dell *et al.* (2014) conclude their review on automated video analysis in ecology with a call to developers. They ask for video analysis systems that are easy to use, do not require marking of individuals, are flexible to work with in a variety of experimental settings and with different organisms, allow tracking of a large number of individuals simultaneously, overcome significant data management issues, and are mostly automated. We believe bemovi is a promising step in this direction and will allow microbial biologists, to follow new and exciting research lines such as the effects of intraspecific and inter-individual variation for ecological and evolutionary dynamics. Furthermore, though we tested utility for microbes, it is likely that bemovi will be useful for analysing any objects moving against a relatively stationary background. For example, insects or birds on a surface could be tracked and analysed, and it might even be possible to track and count birds flying in the sky or fish in the sea (though movement data would need to be treated with caution, given the 2D constraint of bemovi.)

## 5 Acknowledgements

We thank Shawn Vader (uk.linkedin.com/in/shawnvader) for creating the standalone ParticleLinker and the MOSAIC group (http://mosaic.mpi-cbg.de/?q=welcome) for developing the ParticleTracker plug-in in ImageJ. NS is Research Associate of the F.R.S.-FNRS, and acknowledges financial support from ARC 10-15/031, F.R.S.-FNRS and UCL-FSR. FP and OP have financial support by the University of Zurich and Swiss National Science Foundation Grant 31003A_137921 (to OP).

